## Supplementary FIgures for "Unprecedented female mutation bias in aye-ayes"

### Supplementary Figure 1

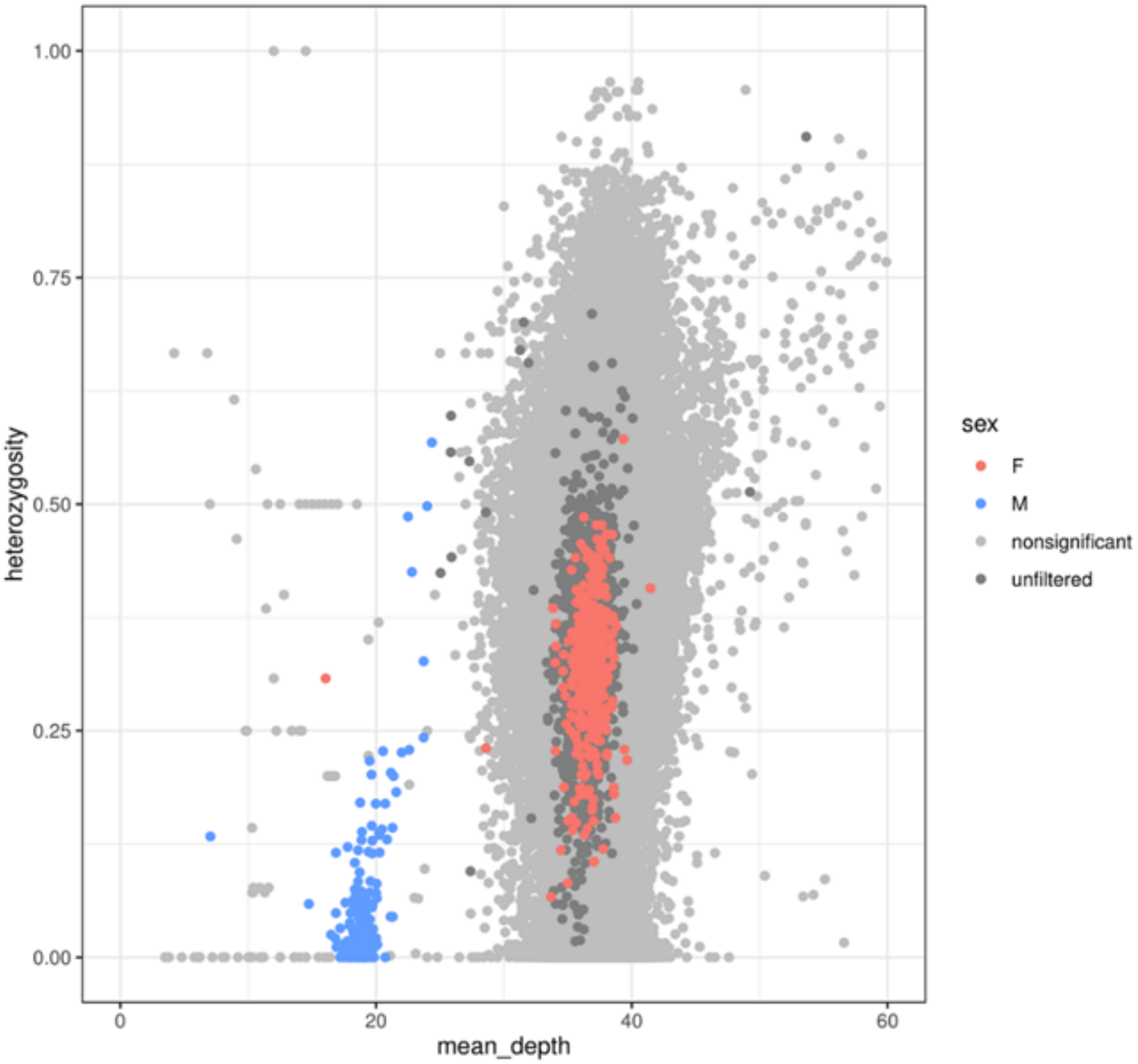

Supplementary Figure 2

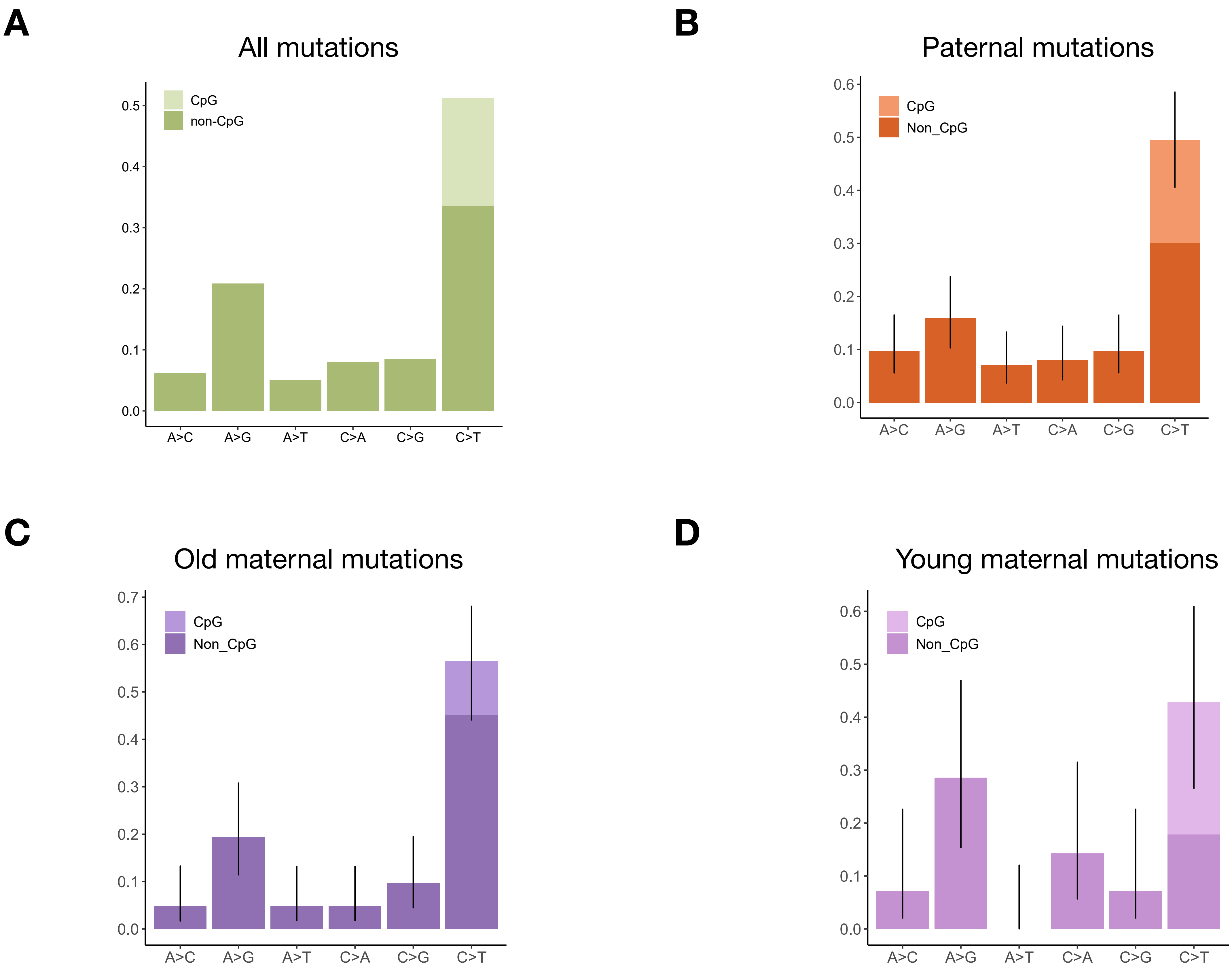

Supplementary Figure 3

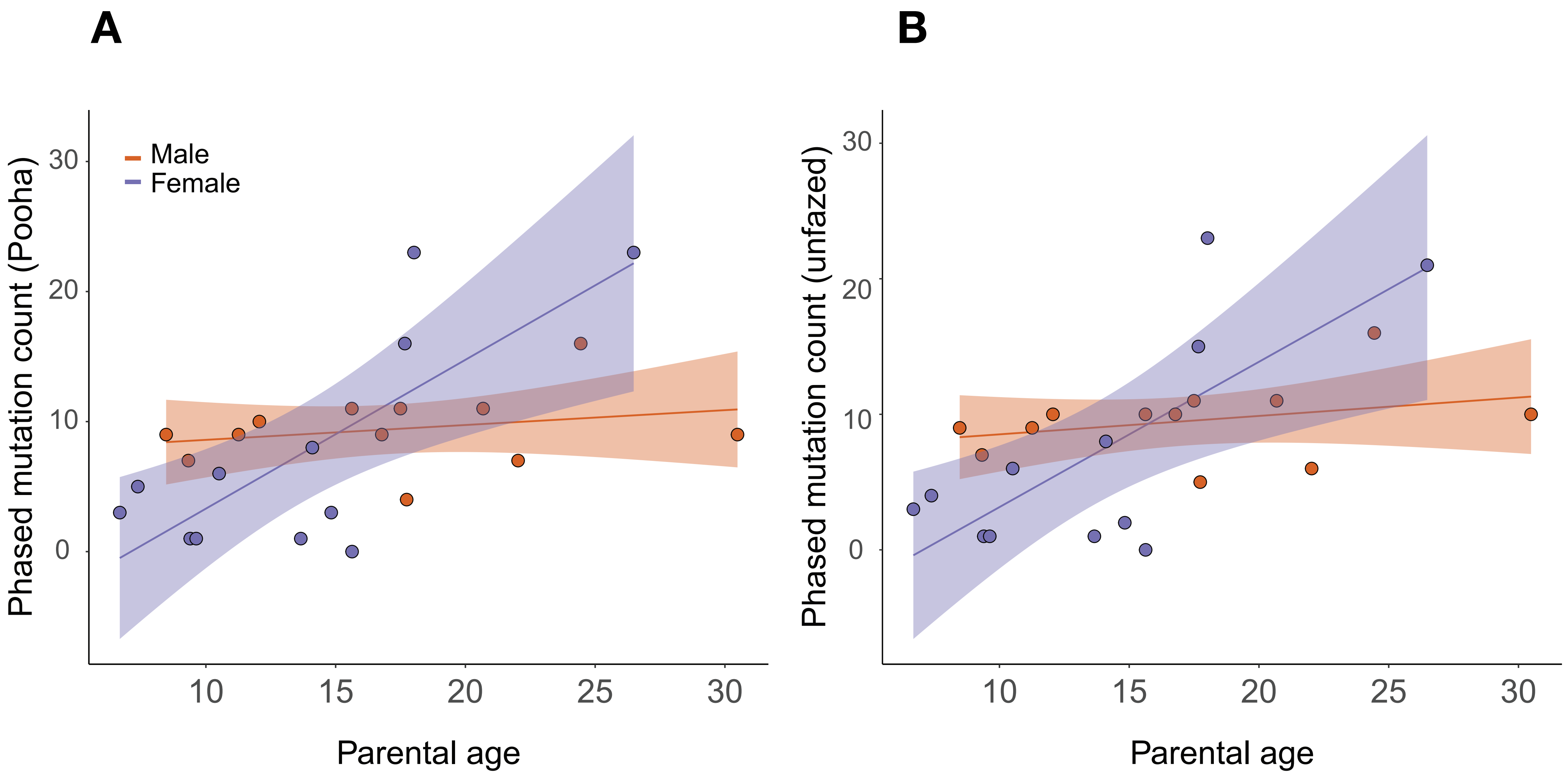

Supplementary Figure 4

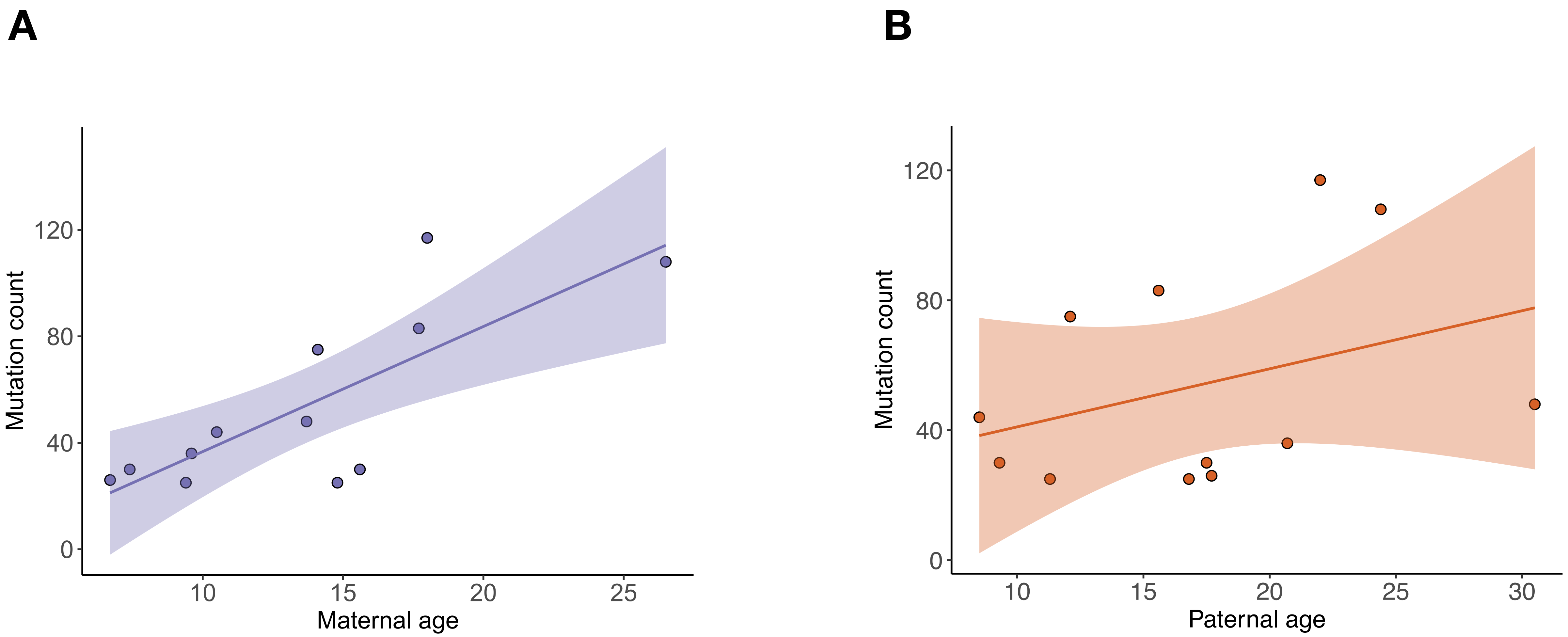

### Supplementary Figure 5

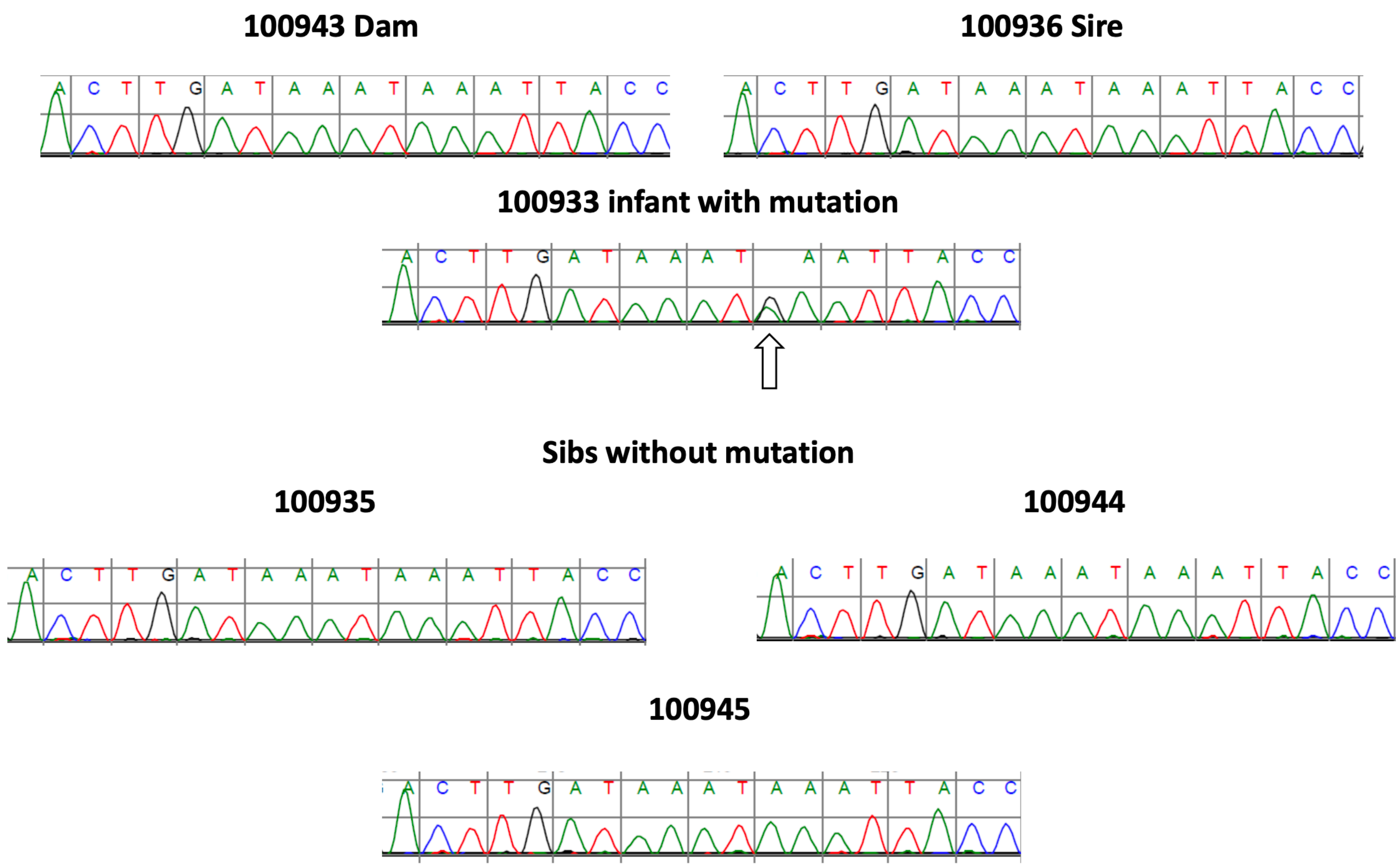

### Supplementary Figure 6

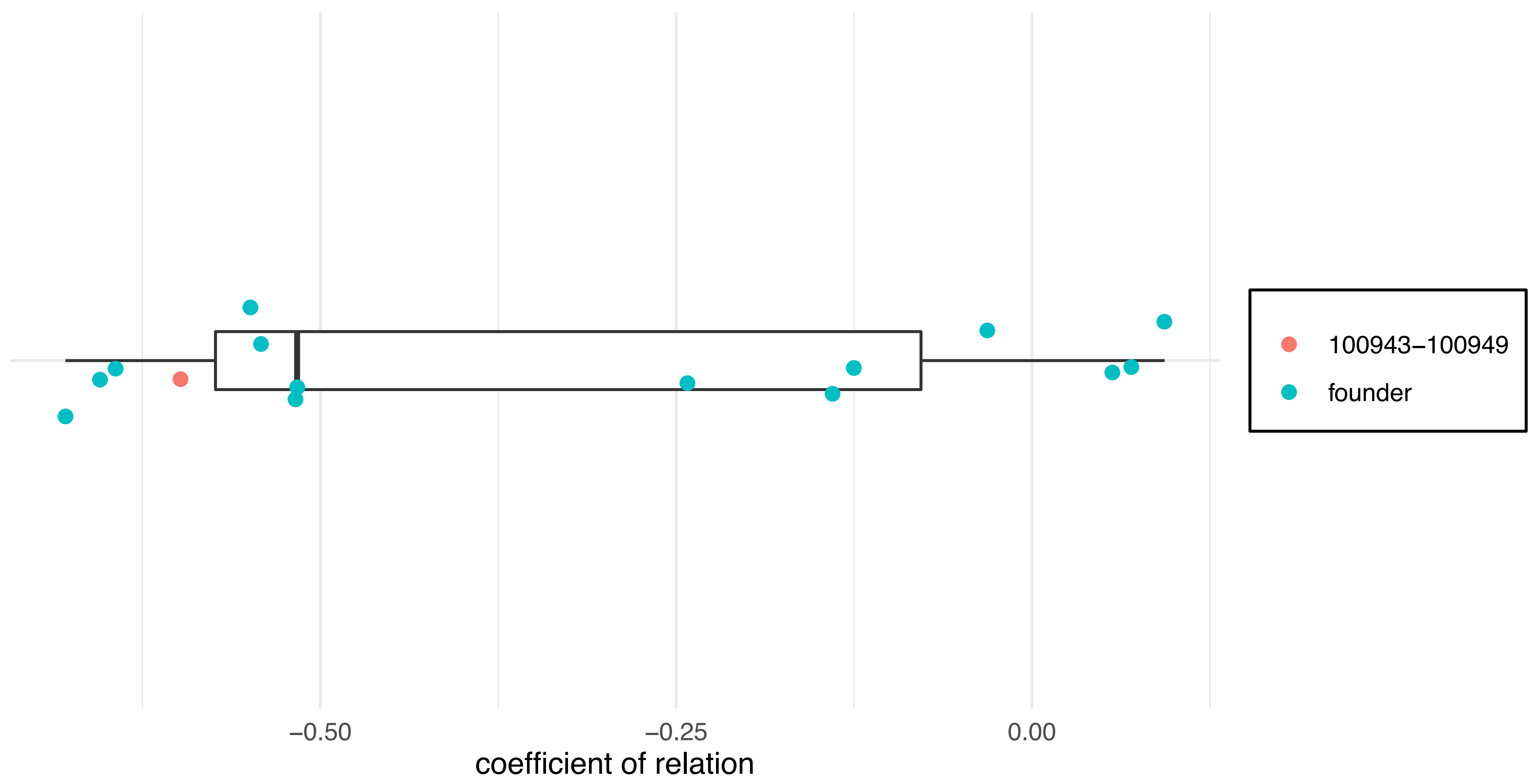

Supplementary Figure 7

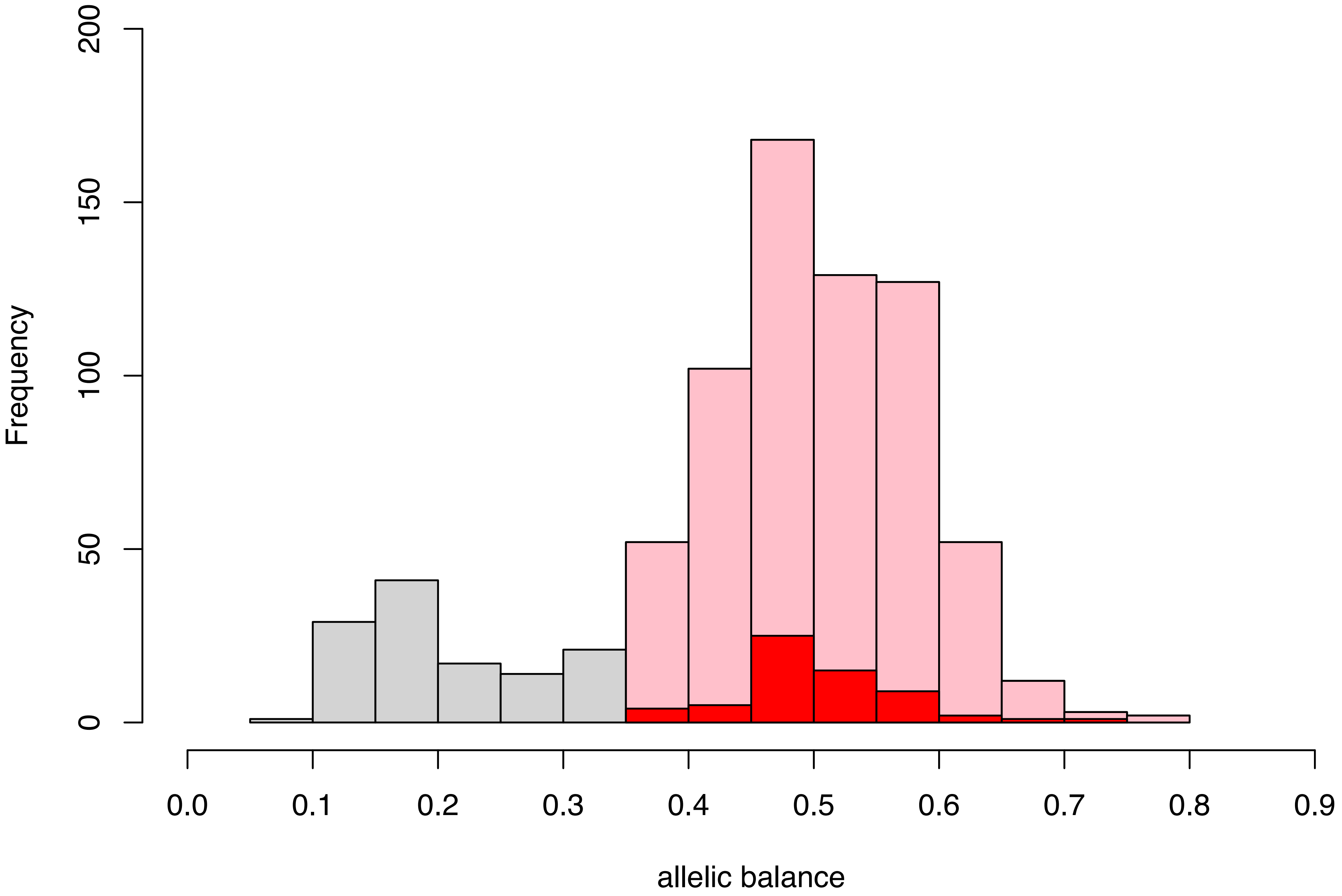

### Supplementary Figure 8

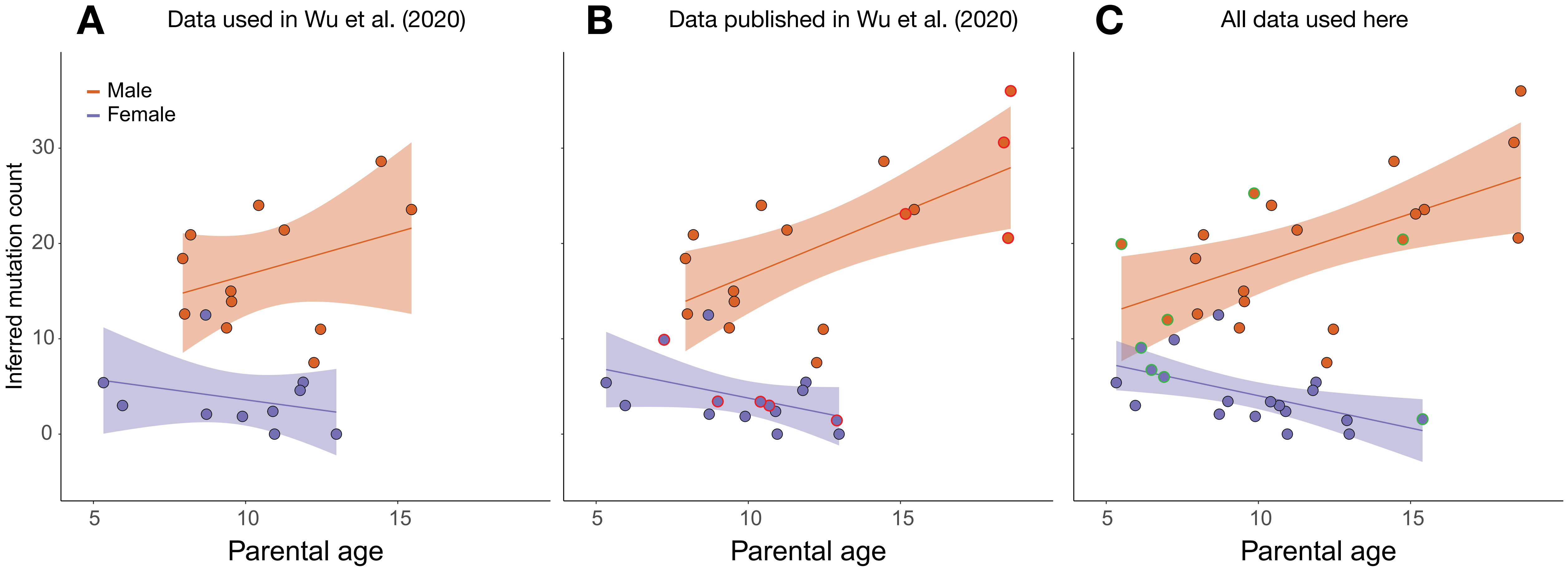
